# ProteoPy: an AnnData-based framework for integrated proteomics analysis

**DOI:** 10.64898/2026.03.31.715273

**Authors:** Ian Dirk Fichtner, Levente Temesvari-Nagy, Felix Sahm, Moritz Gerstung, Isabell Bludau

## Abstract

**Summary:** ProteoPy is a lightweight Python library for protein- and peptide-level quantitative proteomics analysis, built around the AnnData class as its core data structure. It streamlines data import, preprocessing, and differential analysis while preserving all metadata within a single object. A reimplementation of our previously published COPF algorithm enables proteoform group inference directly from peptide-level data, facilitating the identification of proteoform-specific regulation and isoform usage. Designed for accessibility and flexibility, ProteoPy simplifies analysis for non-specialists and provides an extensible foundation for advanced proteomics workflows, seamlessly integrating with the scanpy and muon ecosystems for reproducible and scalable multi-omics analysis.

**Availability and implementation:** ProteoPy is implemented in Python 3 and publicly available on GitHub: https://github.com/UKHD-NP/proteopy under the Apache 2.0 license.

**Supplementary information:** Tutorial notebooks for ProteoPy are included as supplementary data and are also available on GitHub: https://github.com/UKHD-NP/proteopy/tree/main/docs/tutorials.

## Introduction

Mass spectrometry (MS)-based proteomics has become an essential approach for investigating biological systems across a wide range of applications from basic to translational research (Aebersold and Mann, 2003, 2016). Modern workflows enable reproducible quantification of thousands of proteins across large sample cohorts, providing a functional complement to genomic and transcriptomic profiling. These developments are accompanied by a growing interest in integrating proteomics with other molecular modalities to achieve a more comprehensive view of cellular states. At the same time, higher-throughput instrumentation and emerging single-cell and spatial proteomics technologies are generating larger and more complex datasets, bringing analysis requirements closer to those encountered in transcriptomics.

Recent advances in data acquisition and analysis—supported by software such as DIA-NN (Demichev *et al*., 2020), MaxQuant (Cox and Mann, 2008) and AlphaPept (Strauss *et al*., 2024) for data processing, and Perseus (Tyanova *et al*., 2016), MSstats (Kohler *et al*., 2023) or AlphaPeptStats (Krismer *et al*., 2023) for statistical analysis— have made proteomics increasingly robust and quantitative. However, these tools typically rely on distinct data formats and scripting environments, and no widely adopted, unified data structure currently exists. This fragmentation poses several challenges: (i) overlapping functionality is repeatedly implemented across tools, often without leveraging well-established frameworks; (ii) researchers must learn and maintain multiple analysis ecosystems, increasing the barrier to correct and reproducible use; and (iii) integration across omics layers remains cumbersome without a shared data model supporting direct interoperability. In contrast, single-cell and spatial transcriptomics have coalesced around the AnnData (Virshup *et al*., 2024) and scanpy ecosystem (Wolf *et al*., 2018), which provides standardized, scalable, and well-documented pipelines for data handling and analysis—spanning single-cell transcriptomics (scanpy), spatial omics (squidpy) (Palla *et al*., 2022), and multi-omics integration through a shared data format and analysis framework (MuData and muon) (Bredikhin *et al*., 2022). Adopting this existing framework for proteomics would leverage well-established functionality for data processing, visualization, and statistical analysis, enable a seamless transition for researchers familiar with transcriptomic workflows, and facilitate data integration across omics modalities— supporting classical bulk proteomics but also the emerging fields of single-cell and spatially resolved proteomics within a unified computational ecosystem.

At the same time, there is growing recognition that peptide-level analyses can reveal hidden biological information beyond conventional protein-level summaries, enabling the identification of proteoform-specific regulation and isoform usage. Our previously published COPF algorithm illustrates the potential of leveraging peptide-level quantitative patterns to infer proteoform groups across large datasets (Bludau et al., 2021). However, its niche-focused implementation has limited broader adoption until now, underscoring the need for integration into general proteomics workflows.

Here, we present *ProteoPy*, a lightweight Python library that extends the use of the AnnData framework to bulk proteomics at the protein and peptide level. It includes a clean, user-friendly implementation of proteoform group inference and lays the foundation for seamless expansion towards single-cell and spatially resolved proteomics.

## Implementation

### 2.1 Library design

ProteoPy is implemented in Python (≥3.10) and builds on the the AnnData class. Its functions leverage established open-source scientific libraries, including NumPy (Harris et al., 2020), SciPy (Virtanen *et al*., 2020), scikit-learn (Pedregosa *et al*., 2011) and pandas (McKinney, 2010) for omics data handling, and matplotlib (Hunter, 2007) and seaborn (Waskom, 2021) for data visualization. This foundation ensures computational efficiency, reliable numerical operations, and compatibility with widely used analysis workflows. By leveraging the AnnData class, ProteoPy inherits a proven structure for storing quantitative matrices alongside rich metadata. Furthermore, ProteoPy is compatible with external AnnData-based libraries such as scanpy and muon from the scverse ecosystem, enabling seamless interoperability with established single-cell and multi-omics analysis workflows. In particular, integration with muon allows proteomics data to be combined with other omic layers within a unified MuData container, facilitating coordinated preprocessing, joint analysis, and cross-modal exploration in a consistent multi-omics framework.

Function names and syntax in ProteoPy follow the conventions of the scanpy API, including the preprocessing (pp), tools (tl), and plotting (pl) module namespaces, providing a familiar and intuitive interface for users of the *scanpy* ecosystem. The overall structure and functionality of ProteoPy is visualized in Figure 1.

**Figure 1.**
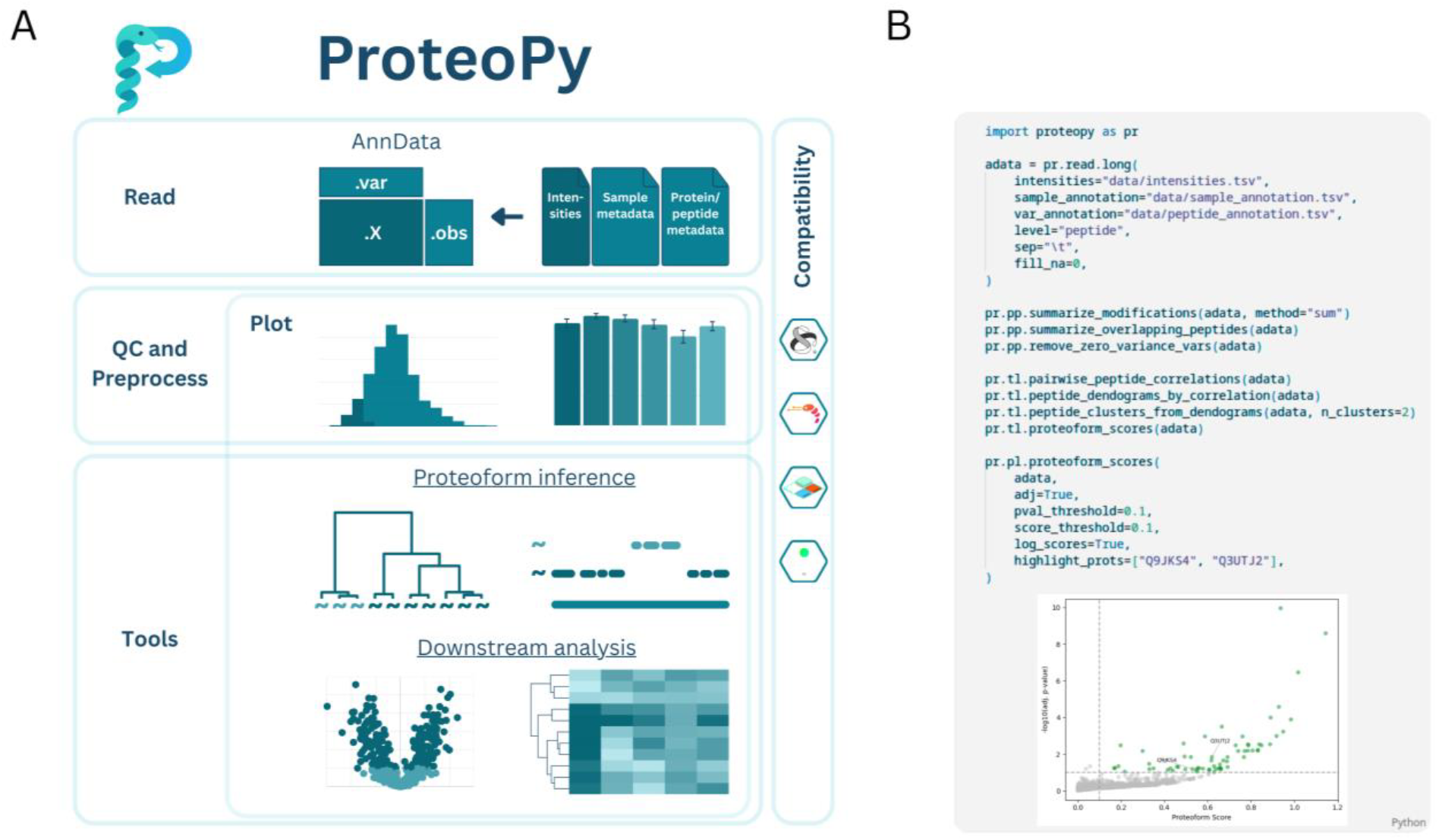
ProteoPy functionality and API design. **A**. The API is organized into four main functional categories. (1) read enables seamless import of data from diverse proteomics acquisition software into the AnnData format. (2) pp (preprocessing) provides functions for quality control and data preprocessing. (3) tl (tools) supports proteoform inference from peptide-level proteomics data and downstream analyses such as differential abundance testing and clustering. (4) plot offers visualization utilities for all analysis steps, facilitating data exploration and interpretation of proteomic results. Built on the AnnData framework, ProteoPy is fully compatible with external AnnData-based packages such as scanpy and muon from the scverse ecosystem. Scverse, scanpy, mudata and muon are trademarks of NumFOCUS. **B**. Example proteoform inference workflow illustrating the concise and intuitive ProteoPy API. The modular read, pp, tl, and pl functions enable streamlined end-to-end analysis pipelines from raw peptide-level data to interpretable proteoform scores and downstream visualizations.

### 2.2 Data import (read)

ProteoPy supports quantitative data import at the protein or peptide level from tools such as DIA-NN (Demichev *et al*., 2020) and other tabular formats. Sample-level annotations, including clinical, experimental, or batch information, can be merged directly during import. Similarly, peptides or proteins can be annotated with metainformation such as protein or gene names and respective gene ontology categories. An AnnData object is constructed from the input data and metainformation and serves as the core data structure for all ProteoPy functions.

### 2.3 Quality control and preprocessing (pp)

ProteoPy includes common routines for quantitative and completeness-based quality control and preprocessing at both the sample (.obs) and feature (.var) level. Summary metrics such as the number and fraction of quantified proteins or peptides per sample, as well as missing-value distributions, intensity ranges and coefficients of variations across sample groups, can be evaluated. Samples or features can then be filtered according to user-defined thresholds for coverage or variability to remove low-quality entries and to ensure consistent data quality. ProteoPy further provides standardized routines for normalization, batch correction, and missing-value imputation. Median normalization is implemented as the default approach to correct for global intensity differences across samples while preserving relative abundance patterns. Batch effects can be addressed directly by leveraging the scanpy ecosystem, for example by using the ComBat algorithm (Johnson *et al*., 2007). ProteoPy includes functionality for missing value imputation using a downshifted Gaussian distribution, following the approach popularized by Perseus (Tyanova *et al*., 2016), where random values are drawn from a normal distribution shifted toward lower intensities. All preprocessing steps can be stored as separate layers in AnnData to ensure full transparency and reversibility.

### 2.4 Tools (tl)

#### 2.4.1 Proteoform inference

One of ProteoPy’s distinctive features is the ability to load peptide-level data to perform proteoform group inference via a Python reimplementation of our previously published COPF algorithm (Bludau *et al*., 2021). This approach exploits peptide-level covariation across samples to identify distinct proteoform patterns. Whereas our original R implementation was primarily designed for protein co-factionation study designs, making its usability for standard proteomics experiments less straight forward, the ProteoPy implementation is flexible and easy to apply across a wide range of experimental setups. Importantly, reliable covariation-based inference requires sufficiently large and heterogeneous datasets comprising multiple biological conditions.

#### 2.4.2 Downstream analysis

ProteoPy supports a broad range of downstream analyses, from unsupervised data exploration to classical differential abundance testing. These analyses are implemented within ProteoPy while maintaining compatibility with external tools through the AnnData data structure. For exploratory analysis, ProteoPy provides functionality for unsupervised clustering. For statistical inference, we implemented a flexible framework for group-wise comparisons, including two-sample t-tests, Welch’s t-test, and one-way ANOVA, together with multiple testing correction using Bonferroni or Benjamini-Hochberg procedures. All results—such as fold changes, p-values, and adjusted p-values—are stored directly within the AnnData object to ensure reproducibility and interoperability. In addition to ProteoPy internal functions, scanpy can also directly be used for various operations such as dimensionality reduction and clustering.

### 2.5 Plotting (pl)

Finally, ProteoPy provides comprehensive visualization capabilities spanning all stages of the analysis workflow, including quality control, preprocessing, proteoform inference, and downstream analyses. The generated plots are designed to be publication-ready, enabling a clear, high-quality presentation of results without extensive post-processing.

## Application of ProteoPy

We applied ProteoPy to two representative MS-based proteomics datasets to demonstrate its capabilities at both the protein and peptide levels. For a general protein-level workflow, we reanalyzed a recent study of human erythropoiesis from Karayel et al. (Karayel *et al*., 2020), reproducing the complete processing pipeline from Spectronaut output to quality control, normalization, imputation, and differential analysis. To highlight peptide-level functionality and proteoform inference, we reanalyzed the mouse tissue dataset from our original COPF study (Bludau *et al*., 2021), recapitulating results within a low-barrier and reproducible framework. All analyses were performed using ProteoPy version 0.1.1, with corresponding Jupyter notebooks provided as Supplementary Notebooks S1–S2 and on the ProteoPy GitHub repository.

## Conclusion and outlook

ProteoPy introduces a unified and extensible framework for quantitative proteomics analysis based on the established AnnData ecosystem. By standardizing data structures and integrating essential preprocessing and statistical workflows, it lowers the barrier to reproducible and accessible proteomics analysis. The inclusion of peptide-level functionality and proteoform inference expands analytical depth beyond conventional protein summarization, enabling more detailed characterization of molecular regulation and diversity. By aligning proteomics with the well-established single-cell and spatial analysis ecosystem in Python, it provides a foundation for future extensions toward single-cell, spatially resolved, and multi-omic proteomics within a shared computational environment.

## Supporting information

Supplementary Notebook S1

Supplementary Notebook S2

## Author contributions

Ian Dirk Fichtner (Conceptualization [equal], Software [lead], Formal analysis [equal], Writing—original draft [supporting], Writing—review & editing [supporting]), Levente Temesvari-Nagy (Software [supporting], Writing—review & editing [supporting]), Felix Sahm (Supervision [supporting], Writing—review & editing [supporting]), Moritz Gerstung (Supervision [supporting], Writing—review & editing [supporting]), and Isabell Bludau (Conceptualization [equal], Software [supporting], Formal analysis [equal], Supervision [lead], Writing—original draft [lead], Writing—review & editing [lead])

## Acknowledgements

The authors thank all members of the Neuropathology Department, especially Armin Hadzic and Dennis Friedel, as well as the Gerstung lab for their continuous support and feedback throughout the completion of the work.

## References

Aebersold, R, and Mann, M (2003). Mass spectrometry-based proteomics. Nature 422, 198–207.

Aebersold, R, and Mann, M (2016). Mass-spectrometric exploration of proteome structure and function. Nature 537, 347–355.

Bludau, I, Frank, M, Dörig, C, Cai, Y, Heusel, M, Rosenberger, G, Picotti, P, Collins, BC, Röst, H, and Aebersold, R (2021). Systematic detection of functional proteoform groups from bottom-up proteomic datasets. Nat Commun 12, 3810.

Bredikhin, D, Kats, I, and Stegle, O (2022). MUON: multimodal omics analysis framework. Genome Biol 23, 1–12.

Cox, J, and Mann, M (2008). MaxQuant enables high peptide identification rates, individualized p.p.b.-range mass accuracies and proteome-wide protein quantification. Nat Biotechnol 26, 1367–1372.

Demichev, V, Messner, CB, Vernardis, SI, Lilley, KS, and Ralser, M (2020). DIA-NN: neural networks and interference correction enable deep proteome coverage in high throughput. Nat Methods 17, 41–44.

Harris, CR, Millman, KJ, Van Der Walt, SJ, Gommers, R, Virtanen, P, Cournapeau, D, Wieser, E, Taylor, J, Berg, S, Smith, NJ, et al. (2020). Array programming with NumPy. Nature 585, 357–362.

Hunter, JD (2007). Matplotlib: A 2D Graphics Environment. Comput Sci Eng 9, 90–95.

Johnson, WE, Li, C, and Rabinovic, A (2007). Adjusting batch effects in microarray expression data using empirical Bayes methods. Biostatistics 8, 118–127.

Karayel, Ö, Xu, P, Bludau, I, Velan Bhoopalan, S. Yao, Y, Ana Rita, FC, Santos, A, Schulman, BA, Alpi, AF, Weiss, MJ, et al. (2020). Integrative proteomics reveals principles of dynamic phosphosignaling networks in human erythropoiesis. Mol Syst Biol 16, MSB20209813.

Kohler, D, Staniak, M, Tsai, T-H, Huang, T, Shulman, N, Bernhardt, OM, MacLean, BX, Nesvizhskii, AI, Reiter, L, Sabido, E, et al. (2023). MSstats Version 4.0:Statistical Analyses of Quantitative Mass Spectrometry-Based Proteomic Experiments with Chromatography-Based Quantification at Scale. J Proteome Res 22, 1466–1482.

Krismer, E, Bludau, I, Strauss, MT, and Mann, M (2023). AlphaPeptStats: an open-source Python package for automated and scalable statistical analysis of mass spectrometry-based proteomics. Bioinformatics 39, btad461.

McKinney, W(2010). Data Structures for Statistical Computing in Python. Austin, Texas: SciPy 2010, 56–61.

Palla, G, Spitzer, H, Klein, M, Fischer, D, Schaar, AC, Kuemmerle, LB, Rybakov, S, Ibarra, IL, Holmberg, O, Virshup, I, et al. (2022). Squidpy: a scalable framework for spatial omics analysis. Nat Methods 19, 171–178.

Pedregosa, F, Varoquaux, G, Gramfort, A, Michel, V, Thirion, B, Grisel, O, Blondel, M, Prettenhofer, P, Weiss, R, Dubourg, V, et al. (2011). Scikit-learn: Machine Learning in Python. Journal of Machine Learning Research 12, 2825–2830.

Strauss, MT, Bludau, I, Zeng, W-F, Voytik, E, Ammar, C, Schessner, JP, Ilango, R, Gill, M, Meier, F, Willems, S, et al. (2024). AlphaPept: a modern and open framework for MS-based proteomics. Nat Commun 15, 2168.

Tyanova, S, Temu, T, Sinitcyn, P, Carlson, A, Hein, MY, Geiger, T, Mann, M, and Cox, J (2016). The Perseus computational platform for comprehensive analysis of (prote)omics data. Nat Methods 13, 731–740.

Virshup, I, Rybakov, S, Theis, FJ, Angerer, P, and Wolf, FA (2024). anndata: Access and store annotated datamatrices. JOSS 9, 4371.

Virtanen, P, Gommers, R, Oliphant, TE, Haberland, M, Reddy, T, Cournapeau, D, Burovski, E, Peterson, P, Weckesser, W, Bright, J, et al. (2020). SciPy 1.0: fundamental algorithms for scientific computing in Python. Nat Methods 17, 261–272.

Waskom, M (2021). seaborn: statistical data visualization. JOSS 6, 3021.

Wolf, FA, Angerer, P, and Theis, FJ (2018). SCANPY: large-scale single-cell gene expression data analysis. Genome Biol 19, 15.

