## Supplementary Notebook S1 for "ProteoPy: an AnnData-based framework for integrated proteomics analysis": supplementary_notebook_S1_protein_analysis.html

karayel-2020\_proteome-remodeling-during-human-erythropoiesis


### Proteome remodeling during human erythropoiesis¶

This notebook demonstrates a comprehensive protein-level analysis workflow using ProteoPy.
To this aim we selected a human erythropoiesis DIA dataset by Karayel et al. (2020):

> Karayel Ö, Xu P, Bludau I, Velan Bhoopalan S, Yao Y, Ana Rita FC, Santos A, Schulman BA, Alpi AF, Weiss MJ, and Mann M. **Integrative proteomics reveals principles of dynamic phosphosignaling networks in human erythropoiesis.** *Molecular Systems Biology*, 16(12):MSB20209813, 2020. doi:10.15252/msb.20209813

The authors profiled the proteome across five stages of *in vitro* erythropoiesis from CD34⁺ hematopoietic stem/progenitor cells (HSPCs) — progenitors, proerythroblasts and early basophilic erythroblasts (ProE&EBaso), late basophilic erythroblasts (LBaso), polychromatic erythroblasts (Poly), and orthochromatic erythroblasts (Ortho) — using DIA-based mass spectrometry with four biological replicates per stage.

Here, we use this dataset to show how ProteoPy can be used to perform typical protein-level analysis steps: quality control, preprocessing, exploratory analysis and differential abundance testing.

#### Setup¶

In [1]:

```
from pathlib import Path
import numpy as np
import scanpy as sc
import matplotlib.pyplot as plt
import matplotlib as mpl
from matplotlib.pyplot import rc_context

import proteopy as pr

# Create a data directory in your current working directory 
# to store files downloaded in this notebook.
cwd = Path(".").resolve()
(cwd / "data").mkdir(parents=True, exist_ok=True)
```

#### Reading in the data¶

The Karayel et al. (2020) dataset contains protein-level DIA-MS intensities from five erythroid differentiation stages of CD34⁺ HSPCs. ProteoPy provides a built-in download function (`pr.download.karayel_2020()`) that fetches the dataset from the PRIDE archive (PXD017276), processes it, and saves it as three separate files that are compatible with the `pr.read.long()` function in ProteoPy:

- **Intensities** — long-format table with columns `sample_id`, `protein_id`, and `intensity` (one row per measurement)
- **Sample annotation** — maps each `sample_id` to its `cell_type` and `replicate`
- **Protein annotation** — maps each `protein_id` to its `gene_id`

In [2]:

```
# Define paths to the data files. These are the same paths that will be used in the download function below.
intensities_path = (
    "data/karayel-2020_ms-proteomics"
    "_human-erythropoiesis_intensities.tsv"
)
sample_annotation_path = (
    "data/karayel-2020_ms-proteomics"
    "_human-erythropoiesis_sample-annotation.tsv"
)
var_annotation_path = (
    "data/karayel-2020_ms-proteomics"
    "_human-erythropoiesis_protein-annotation.tsv"
)

pr.download.karayel_2020(
    intensities_path=intensities_path,
    sample_annotation_path=sample_annotation_path,
    var_annotation_path=var_annotation_path,
    force=True,
)
```

A quick look at the first rows of each file shows the data format ProteoPy expects for each information layer:

In [3]:

```
# intensities
!head -n5 {intensities_path}
```

```
sample_id	protein_id	intensity
LBaso_rep1	A0A024QZ33;Q9H0G5	15084.2626953125
LBaso_rep2	A0A024QZ33;Q9H0G5	15442.2646484375
LBaso_rep3	A0A024QZ33;Q9H0G5	13992.021484375
LBaso_rep4	A0A024QZ33;Q9H0G5	18426.5390625
```

In [4]:

```
# sample_annotation
!head -n5 {sample_annotation_path}
```

```
sample_id	cell_type	replicate
LBaso_rep1	LBaso	rep1
LBaso_rep2	LBaso	rep2
LBaso_rep3	LBaso	rep3
LBaso_rep4	LBaso	rep4
```

In [5]:

```
# var_annotation
!head -n5 {var_annotation_path}
```

```
protein_id	gene_id
A0A024QZ33;Q9H0G5	NSRP1
A0A024QZB8;A0A0A0MRC7;A0A0D9SFH9;A0A1B0GV71;A0A1B0GW34;A0A1B0GWD3;H3BR00;Q13286;Q13286-2;Q13286-3;Q13286-4;Q13286-5;Q9UBD8	CLN3;CLN3;CLN3;CLN3;CLN3;;CLN3;CLN3;CLN3;CLN3;CLN3;CLN3;CLN3
A0A024R0Y4;O75478	TADA2A
A0A024R216;Q9Y3E1	HDGFRP3;HDGFL3
```

With the three files on disk, `pr.read.long()` reads the long-format intensities, pivots them into a dense matrix, and merges the sample and protein annotations into a single AnnData object. Setting `fill_na=0` replaces any missing intensity values with zero.

In [6]:

```
# Create a protein level AnnData object from the 
# downloaded data files and fill missing values with 0.
adata = pr.read.long(
    intensities=intensities_path,
    sample_annotation=sample_annotation_path,
    var_annotation=var_annotation_path,
    level="protein",
    fill_na=0,
)

adata
```

Out[6]:

```
AnnData object with n_obs × n_vars = 20 × 7758
    obs: 'sample_id', 'cell_type', 'replicate'
    var: 'protein_id', 'gene_id'
```

In [7]:

```
# Check that the created AnnData object is formatted correctly following
# proteodata conventions defined in ProteoPy
print("Is proteodata: ", pr.utils.is_proteodata(adata))
```

```
Is proteodata:  (True, 'protein')
```

You can define consistent color schemes and category orders for the differentiation stages and replicates used for all figures created by ProteoPy.

In [8]:

```
# Cell types
adata.uns["colors_cell_type"] = {
    "LBaso": "#D91C5C",
    "Ortho": "#F6922F",
    "Poly": "#262261",
    "ProE&EBaso": "#1B75BB",
    "Progenitor": "#D6DE3B",
}

adata.uns["order_cell_type"] = ["Progenitor", "ProE&EBaso", "LBaso", "Poly", "Ortho"]

# Replicates
n_reps = adata.obs["replicate"].nunique()
cmap = mpl.colormaps["tab10"]
adata.uns["colors_replicate"] = cmap(range(n_reps)).tolist()

adata.uns["order_replicate"] = sorted(adata.obs["replicate"].unique().tolist())
```

Looking at the sample distribution across categories shows that the data was successfully imported, including a total of 20 samples derived from 5 differentiation stages with 4 biological replicates each.

In [9]:

```
pr.pl.n_samples_per_category(
    adata,
    category_key="cell_type",
    color_scheme=adata.uns["colors_cell_type"],
    order=adata.uns["order_cell_type"],
)
```

In [10]:

```
pr.pl.n_samples_per_category(
    adata,
    category_key="replicate",
    color_scheme=adata.uns["colors_replicate"],
    order=adata.uns["order_replicate"],
)
```

#### Quality control and preprocessing¶

##### Detected proteins per sample¶

In a first QC step, you can check the number of proteins detected in each sample. The per-sample detected proteins plot shows a decrease along the differentiation trajectory (left -> right order).

In [11]:

```
pr.pl.n_proteins_per_sample(
    adata,
    zero_to_na=True,
    order_by="cell_type",
    order=adata.uns["order_cell_type"],
    color_scheme=adata.uns["colors_cell_type"],
    xlabel_rotation=45,
    print_stats=True,
)
```

```
Global:
 mean_count  std_count  median_count  min_count  max_count  mean_pct  std_pct  median_pct  min_pct  max_pct
     6925.6      623.0        7160.5       5490       7565      89.3      8.0        92.3     70.8     97.5

Per cell_type:
 cell_type  mean_count  std_count  median_count  min_count  max_count  mean_pct  std_pct  median_pct  min_pct  max_pct
     LBaso      7165.2      150.2        7160.5       7027       7313      92.4      1.9        92.3     90.6     94.3
     Ortho      5866.5      257.5        5952.0       5490       6072      75.6      3.3        76.7     70.8     78.3
      Poly      6738.8      232.7        6778.0       6426       6973      86.9      3.0        87.4     82.8     89.9
ProE&EBaso      7369.8       68.8        7365.5       7290       7458      95.0      0.9        94.9     94.0     96.1
Progenitor      7487.5       88.7        7512.5       7360       7565      96.5      1.1        96.8     94.9     97.5
```

Out[11]:

```
<Axes: ylabel='#'>
```

##### Completeness filtering¶

To ensure reliable downstream analysis, proteins that are only sporadically measured in a proteomics experiment can be removed using a completeness filtering function. In this dataset, proteins are retained if they are detected in all replicates of at least one differentiation stage, ensuring that only consistently quantified proteins remain.

In [12]:

```
pr.pp.filter_var_completeness(
    adata,
    min_fraction=1,
    group_by="cell_type",
    zero_to_na=True,
)
```

```
279 var removed
```

In [13]:

```
pr.pl.completeness_per_var(adata, zero_to_na=True)
```

Out[13]:

```
<Axes: title={'center': 'Completeness per var'}, xlabel='Fraction of non-missing obs values per var', ylabel='Count'>
```

In [14]:

```
print(
    f"{adata.n_vars} protein groups remain after "
    f"filtering for group-wise completeness"
)
```

```
7479 protein groups remain after filtering for group-wise completeness
```

##### Detected proteins per sample¶

After filtering, 7,479 protein groups remain.

In [15]:

```
# Group samples by cell type
pr.pl.n_proteins_per_sample(
    adata,
    zero_to_na=True,
    group_by="cell_type",
    order=adata.uns["order_cell_type"],
    color_scheme=adata.uns["colors_cell_type"],
    xlabel_rotation=45,
)
```

Out[15]:

```
<Axes: xlabel='cell_type', ylabel='#'>
```

##### Abundance rank plot¶

Abundance rank plots show the dynamic range and ranking of marker proteins across stages. Intensity signals span five (Progenitor) to seven (Ortho) orders of magnitude. As expected, Haemoglobin subunits generally rank among the highest expressed proteins and increase by approximately three orders of magnitude from the Progenitor to Ortho stage.

In [16]:

```
haemoglobin = {
    "P68871": "HBB",
    "P69905": "HBA1",
    "P02100": "HBE1",
    "A0A2R8Y7X9;P69891": "HBG1",
}

pr.pl.abundance_rank(
    adata,
    color="cell_type",
    log_transform=10,
    zero_to_na=True,
    color_scheme=adata.uns["colors_cell_type"],
    summary_method="median",
    highlight_vars=list(haemoglobin.keys()),
    var_labels_key="gene_id",
)
```

##### Coefficient of variation¶

Coefficients of variation (CVs) can be checked to inspect overall data consistency. Here, intra-stage coefficients of variation (CVs) confirm high reproducibility across replicates of each differentiation stage.

In [17]:

```
pr.pl.cv_by_group(
    adata,
    group_by="cell_type",
    min_samples=3,
    zero_to_na=True,
    color_scheme=adata.uns["colors_cell_type"],
    figsize=(6, 4),
    xlabel_rotation=45,
    order=adata.uns["order_cell_type"],
    hline=0.2,
    print_stats=True,
)
```

```
Global CV Summary:
 Count    Min   Max  Median   Mean    Std
 34054 0.0021 1.971  0.1099 0.1725 0.1887

Per-Group CV Summary:
            Count     Min     Max  Median    Mean     Std
Group                                                    
Progenitor   7366  0.0051  1.8955  0.1219  0.1899  0.2054
ProE&EBaso   7261  0.0038  1.8931  0.0774  0.1336  0.1674
LBaso        7058  0.0046  1.8879  0.1147  0.1736  0.1842
Poly         6646  0.0047  1.9710  0.1292  0.1878  0.1887
Ortho        5723  0.0021  1.9081  0.1176  0.1805  0.1904

Global Threshold Summary (hline=0.2):
 Count below  Percentage below
       25308           74.3173

Per-Group Threshold Summary (hline=0.2):
            Count below  Percentage below
Group                                    
Progenitor       5151.0           69.9294
ProE&EBaso       6082.0           83.7626
LBaso            5322.0           75.4038
Poly             4642.0           69.8465
Ortho            4111.0           71.8330
```

##### Log₂ transformation¶

In [18]:

```
# Convert zeros to NaN and log2 transform
adata.layers["raw"] = adata.X.copy()
adata.X[adata.X == 0] = np.nan
adata.X = np.log2(adata.X)
```

##### Median normalization¶

In [19]:

```
# Intensity boxplot before normalization
pr.pl.intensity_box_per_sample(
    adata,
    order_by="cell_type",
    order=adata.uns["order_cell_type"],
    zero_to_na=True,
    color_scheme=adata.uns["colors_cell_type"],
    xlabel_rotation=45,
    figsize=(7, 4),
)
```

Out[19]:

```
<Axes: ylabel='Intensity'>
```

In [20]:

```
pr.pp.normalize_median(adata, log_space=True)
```

In [21]:

```
# Intensity boxplot after normalization: median values across samples are the same
pr.pl.intensity_box_per_sample(
    adata,
    order_by="cell_type",
    order=adata.uns["order_cell_type"],
    zero_to_na=True,
    color_scheme=adata.uns["colors_cell_type"],
    xlabel_rotation=45,
    figsize=(7, 4),
)
```

Out[21]:

```
<Axes: ylabel='Intensity'>
```

##### Missing value imputation¶

Missing values are imputed using a downshifted Gaussian distribution, following the approach popularized by Perseus (Tyanova et al., 2016), where random values are drawn from a normal distribution shifted toward lower intensities. The approach assumes that missing values arise from low-abundance proteins below the detection limit (missing not at random, MNAR).

In [22]:

```
pr.pp.impute_downshift(
    adata,
    downshift=1.8,
    width=0.3,
    zero_to_na=True,
    random_state=123,
    verbose=True,
)
```

```
Measured: 137,154 values (91.7%)
Imputed: 12,426 values (8.3%)
```

Intensity distributions with imputed values highlighted confirm that imputed values form a distinct low-intensity shoulder:

In [23]:

```
# Combined histogram across all samples
pr.pl.intensity_hist(
    adata,
    color_imputed=True,
    density=False,
)

# Per-sample histograms (small multiples)
pr.pl.intensity_hist(
    adata,
    color_imputed=True,
    per_obs=True,
    ncols=5,
    legend_loc="upper right",
    density=False,
    figsize=(17, 12),
)
```

#### Exploratory data analysis¶

##### Sample correlation matrix¶

A sample correlation matrix can be calculated and plotted to inspect which samples share overall expression patterns. Here, high intra-stage (Pearson *r* > 0.94) and lower inter-stage Pearson correlations confirm that protein abundances are primarily driven by differentiation stage.

In [24]:

```
pr.pl.sample_correlation_matrix(
    adata,
    margin_color="cell_type",
    color_scheme=adata.uns["colors_cell_type"],
    xticklabels=True,
    figsize=(9, 8),
    print_stats=True,
)
```

```
Sample correlation summary (off-diagonal, pearson):
     min      max     mean   median      std
0.772086 0.976173 0.901526 0.909919 0.049865

Per-sample mean correlation:
      sample_id  mean_corr
     Ortho_rep4   0.864798
     Ortho_rep3   0.885271
     Ortho_rep1   0.872628
     Ortho_rep2   0.881339
Progenitor_rep4   0.856958
Progenitor_rep3   0.866849
Progenitor_rep1   0.891815
Progenitor_rep2   0.898941
     LBaso_rep1   0.924209
     LBaso_rep2   0.925118
ProE&EBaso_rep4   0.918244
ProE&EBaso_rep1   0.917999
ProE&EBaso_rep2   0.920335
ProE&EBaso_rep3   0.919495
      Poly_rep3   0.911729
      Poly_rep4   0.901124
      Poly_rep1   0.912838
     LBaso_rep4   0.921740
     LBaso_rep3   0.921465
      Poly_rep2   0.917615

Per-group mean correlation:
     group  mean_within  mean_between
     LBaso     0.955621      0.917042
     Ortho     0.941910      0.863653
      Poly     0.950539      0.903380
ProE&EBaso     0.968593      0.909723
Progenitor     0.954879      0.864346
```

##### Dimensionality reduction¶

The AnnData-based proteodata object created and used by ProteoPy is directly compatible with other AnnData based python packages such as scanpy (`import scanpy as sc`), which can for example be used for dimensionality reduction and clustering.

Here, the PCA and UMAP analyses further confirm that differentiation stage is the dominant source of variation in the dataset.

###### PCA analysis¶

In [25]:

```
sc.tl.pca(adata)
with rc_context({"figure.figsize": (5, 3)}):
    sc.pl.pca_variance_ratio(adata, n_pcs=50, log=True)

sc.pl.pca(
    adata,
    color="cell_type",
    dimensions=(0, 1),
    ncols=2,
    size=100,
    palette=adata.uns["colors_cell_type"],
)

sc.pl.pca(
    adata,
    color="replicate",
    dimensions=(0, 1),
    ncols=2,
    size=100,
    palette=adata.uns["colors_replicate"],
)
```

###### UMAP¶

In [26]:

```
sc.pp.neighbors(adata, n_neighbors=3)
sc.tl.umap(adata)

sc.pl.umap(
    adata,
    color="cell_type",
    size=100,
    palette=adata.uns["colors_cell_type"],
)

sc.pl.umap(
    adata,
    color="replicate",
    size=100,
    palette=adata.uns["colors_replicate"],
)
```

#### Differential abundance analysis¶

ProteoPy includes functionality for basic statistical testing across conditions, for example using the Student's t-test and visualization by volcano plots.

In this erythropoiesis dataset, pairwise comparisons of each stage against the progenitor stage via the Student's t-test reveal cumulative proteome remodeling. The later maturation stages (Poly, Ortho) show a larger number of differentially abundant proteins, reflecting the progressive divergence from the progenitor state.

In [27]:

```
progenitor_ct = adata.uns["order_cell_type"][0]
rest_ct = adata.uns["order_cell_type"][1:]
for ct in rest_ct:
    pr.tl.differential_abundance(
        adata,
        method="ttest_two_sample",
        group_by="cell_type",
        setup={"group1": ct, "group2": progenitor_ct},
        multitest_correction="bh",
        alpha=0.01,
        space="log",
    )
```

```
Saved test results in .varm['ttest_two_sample;cell_type;ProE_EBaso_vs_Progenitor']
Saved test results in .varm['ttest_two_sample;cell_type;LBaso_vs_Progenitor']
Saved test results in .varm['ttest_two_sample;cell_type;Poly_vs_Progenitor']
Saved test results in .varm['ttest_two_sample;cell_type;Ortho_vs_Progenitor']
```

Volcano plots with raw and adjusted p-value significance

In [28]:

```
# Volcano plots for progenitor vs subsequent differentiation stages
for ct in rest_ct:
    test_slot = (
        f"ttest_two_sample;cell_type;"
        f"{ct.replace('&', '_')}_vs_Progenitor"
    )

    sig_df = pr.get.differential_abundance_df(
        adata, keys=test_slot,
    )
    sig_series = (
        (sig_df["pval_adj"] <= 0.01)
        & (sig_df["logfc"].abs() >= 0.5)
    )
    print(
        f"{ct} vs Progenitor: \n"
        f"{sum(sig_series)} "
        f"({round(sum(sig_series) / adata.n_vars * 100, 1)}%) "
        f"significant differentially abundant "
        f"protein groups found."
    )

    fig, axes = plt.subplots(ncols=2, figsize=(16, 7))

    # Raw p-values, colored by adjusted significance
    pr.pl.volcano(
        adata,
        varm_slot=test_slot,
        top_labels=5,
        alt_labels_key="gene_id",
        xlabel="log2FC",
        ylabel="pval",
        pval_col="pval",
        alt_color=sig_series,
        title=f"{ct} vs Progenitor",
        ax=axes[0],
        show=False,
    )

    # Adjusted p-values
    pr.pl.volcano(
        adata,
        varm_slot=test_slot,
        fc_thresh=0.5,
        pval_thresh=0.01,
        top_labels=5,
        alt_labels_key="gene_id",
        xlabel="log2FC",
        ylabel="pval_adj",
        title=f"{ct} vs Progenitor",
        ax=axes[1],
        show=False,
    )

    plt.show()
```

```
ProE&EBaso vs Progenitor: 
569 (7.6%) significant differentially abundant protein groups found.
```

```
LBaso vs Progenitor: 
1000 (13.4%) significant differentially abundant protein groups found.
```

```
Poly vs Progenitor: 
2092 (28.0%) significant differentially abundant protein groups found.
```

```
Ortho vs Progenitor: 
3908 (52.3%) significant differentially abundant protein groups found.
```

#### Summary¶

This notebook shows an exemplary protein-level analysis workflow for proteomics
data using ProteoPy. Applied to the erythropoiesis dataset by Karayel et al. (2020),
ProteoPy can reproduce the key data processing steps from quality control,
over preprocessing and exploratory data analysis to differential testing.

In [29]:

```
# Documentation of the used version of proteopy and 
# its dependencies for reproducibility
!pip freeze
```

```
adjustText==1.3.0
anndata==0.11.4
anyio==4.12.1
argon2-cffi==25.1.0
argon2-cffi-bindings==25.1.0
array-api-compat==1.14.0
arrow==1.4.0
asttokens==3.0.1
async-lru==2.3.0
attrs==26.1.0
babel==2.18.0
beautifulsoup4==4.14.3
biopython==1.86
bleach==6.3.0
certifi==2026.2.25
cffi==2.0.0
charset-normalizer==3.4.6
comm==0.2.3
contourpy==1.3.2
cycler==0.12.1
debugpy==1.8.20
decorator==5.2.1
defusedxml==0.7.1
et_xmlfile==2.0.0
exceptiongroup==1.3.1
executing==2.2.1
fastjsonschema==2.21.2
fonttools==4.62.1
fqdn==1.5.1
h11==0.16.0
h5py==3.16.0
httpcore==1.0.9
httpx==0.28.1
idna==3.11
igraph==1.0.0
ipykernel==7.2.0
ipython==8.38.0
isoduration==20.11.0
jedi==0.19.2
Jinja2==3.1.6
joblib==1.5.3
json5==0.13.0
jsonpointer==3.1.1
jsonschema==4.26.0
jsonschema-specifications==2025.9.1
jupyter-events==0.12.0
jupyter-lsp==2.3.0
jupyter_client==8.8.0
jupyter_core==5.9.1
jupyter_server==2.17.0
jupyter_server_terminals==0.5.4
jupyterlab==4.5.6
jupyterlab_pygments==0.3.0
jupyterlab_server==2.28.0
kiwisolver==1.5.0
lark==1.3.1
legacy-api-wrap==1.5
llvmlite==0.46.0
MarkupSafe==3.0.3
matplotlib==3.10.8
matplotlib-inline==0.2.1
mistune==3.2.0
natsort==8.4.0
nbclient==0.10.4
nbconvert==7.17.0
nbformat==5.10.4
nest-asyncio==1.6.0
networkx==3.4.2
notebook_shim==0.2.4
numba==0.64.0
numpy==2.2.6
openpyxl==3.1.5
overrides==7.7.0
packaging @ file:///home/conda/feedstock_root/build_artifacts/bld/rattler-build_packaging_1769093650/work
pandas==2.3.3
pandocfilters==1.5.1
parso==0.8.6
patsy==1.0.2
pexpect==4.9.0
pillow==12.1.1
platformdirs==4.9.4
pooch==1.9.0
prometheus_client==0.24.1
prompt_toolkit==3.0.52
proteopy==0.1.1
psutil==7.2.2
ptyprocess==0.7.0
pure_eval==0.2.3
pyarrow==23.0.1
pycparser==3.0
Pygments==2.19.2
pynndescent==0.6.0
pyparsing==3.3.2
python-dateutil==2.9.0.post0
python-igraph==1.0.0
python-json-logger==4.0.0
pytz==2026.1.post1
PyYAML==6.0.3
pyzmq==27.1.0
referencing==0.37.0
requests==2.32.5
rfc3339-validator==0.1.4
rfc3986-validator==0.1.1
rfc3987-syntax==1.1.0
rpds-py==0.30.0
scanpy==1.11.5
scikit-learn==1.7.2
scipy==1.15.3
seaborn==0.13.2
Send2Trash==2.1.0
session-info2==0.4
six==1.17.0
soupsieve==2.8.3
stack-data==0.6.3
statsmodels==0.14.6
terminado==0.18.1
texttable==1.7.0
threadpoolctl==3.6.0
tinycss2==1.4.0
tomli==2.4.0
tornado==6.5.5
tqdm==4.67.3
traitlets==5.14.3
typing_extensions==4.15.0
tzdata==2025.3
umap-learn==0.5.11
uri-template==1.3.0
urllib3==2.6.3
wcwidth==0.6.0
webcolors==25.10.0
webencodings==0.5.1
websocket-client==1.9.0
```
