## Supplementary Notebook S2 for "ProteoPy: an AnnData-based framework for integrated proteomics analysis": supplementary_notebook_S2_proteoform_inference.html

bludau-2021\_tissue-specific-proteoform-inference-across-five-mouse-organs


### Tissue-specific proteoform inference across five mouse organs¶

This notebook shows how to use ProteoPy for proteoform group inference following a previously established approach that we published in Bludau et al. (2021):

> Bludau I, Frank M, Dörig C, Cai Y, Heusel M, Rosenberger G, Picotti P, Collins BC, Röst H, and Aebersold R. **Systematic detection of functional proteoform groups from bottom-up proteomic datasets.** *Nature Communications*, 12:3810, 2021. doi:10.1038/s41467-021-24030-x

In this paper, we introduced COPF (COrrelation-based functional ProteoForm assessment), a data-driven strategy that assigns peptides to co-varying proteoform groups based on peptide correlation patterns. It was applied to SWATH-MS (DIA) data from five mouse tissues (brain, brown adipose tissue, heart, liver, and quadriceps) across eight BXD mice, originally published by Williams et al. (2018):

> Williams EG, Wu Y, Wolski W, Kim JY, Lan J, Hasan M, Halter C, Jha P, Ryu D, Auwerx J, and Aebersold R. **Quantifying and Localizing the Mitochondrial Proteome Across Five Tissues in A Mouse Population.** *Molecular & Cellular Proteomics*, 17(9):1766–1777, 2018. doi:10.1074/mcp.RA118.000554

In this notebook, we show how the COPF implementation in ProteoPy can be used to perform the core proteoform inference workflow.

#### Setup¶

In [1]:

```
import random
from pathlib import Path
import numpy as np
import scanpy as sc
import matplotlib as mpl
from matplotlib.pyplot import rc_context

import proteopy as pr  # Convention: import proteopy as pr
from proteopy.utils import is_proteodata

# Set random seed for reproducibility
random.seed(42)

# Create a data directory in your current working directory 
# to store files downloaded in this notebook.
cwd = Path(".").resolve()
(cwd / 'data').mkdir(parents=True, exist_ok=True)
```

#### Reading in the data¶

The Williams et al. (2018) dataset contains peptide-level SWATH-MS intensities from five tissues of eight BXD mice. ProteoPy provides a built-in download function (`pr.download.williams_2018()`) that fetches the dataset from the original supplementary archive, processes it, and saves it as three separate files:

- **Intensities** — long-format table with columns `sample_id`, `peptide_id`, and `intensity` (one row per measurement)
- **Sample annotation** — maps each `sample_id` to its `tissue` and `mouse_id`
- **Peptide annotation** — maps each `peptide_id` to its `protein_id` and `gene_id`

This separation into three files mirrors a common data delivery format in proteomics. The files are then read into an AnnData object using `pr.read.long()`.

In [2]:

```
# Define paths to the data files. These are the same paths that will be used in the download function below.
intensities_path = "data/williams-2018_mouse-tissue_intensities.tsv"
sample_annotation_path = "data/williams-2018_mouse-tissue_sample_annotation.tsv"
peptide_annotation_path = "data/williams-2018_mouse-tissue_peptide_annotation.tsv"

pr.download.williams_2018(
    intensities_path=intensities_path,
    sample_annotation_path=sample_annotation_path,
    var_annotation_path=peptide_annotation_path,
    force=True,  # Overwrite files if already exist
)
```

A quick look at the first rows of each file shows the data format ProteoPy expects for each information layer:

In [3]:

```
# intensities
!head -n5 {intensities_path}
```

```
sample_id	peptide_id	intensity
C57_Brain	AAAAAAAAAAAAAAAGAAGK	26302.0
DBA_Brain	AAAAAAAAAAAAAAAGAAGK	15460.0
45_Brain	AAAAAAAAAAAAAAAGAAGK	24631.0
66_Brain	AAAAAAAAAAAAAAAGAAGK	26554.0
```

In [4]:

```
# sample_annotation
!head -n5 {sample_annotation_path}
```

```
sample_id	tissue	mouse_id
C57_Brain	Brain	C57
DBA_Brain	Brain	DBA
45_Brain	Brain	45
66_Brain	Brain	66
```

In [5]:

```
# var_annotation
!head -n5 {peptide_annotation_path}
```

```
peptide_id	protein_id	gene_id
AAAAAAAAAAAAAAAGAAGK	P55012	Slc12a2
AAAAADLANR	Q9JHS4	Clpx
AAAADGEPLHNEEER	Q80WW9	Ddrgk1
AAAAEGARPLER	Q80UM7	Mogs
```

With the three files on disk, `pr.read.long()` reads the long-format intensities, pivots them into a dense matrix, and merges the sample and peptide annotations into a single AnnData object. Setting `fill_na=0` replaces any missing intensity values with zero.

In [6]:

```
# Create a peptide level AnnData object from the 
# downloaded data files and fill missing values with 0.
adata = pr.read.long(
    intensities=intensities_path,
    sample_annotation=sample_annotation_path,
    var_annotation=peptide_annotation_path,
    level="peptide",
    fill_na=0,
)

adata
```

Out[6]:

```
AnnData object with n_obs × n_vars = 40 × 32690
    obs: 'sample_id', 'tissue', 'mouse_id'
    var: 'peptide_id', 'protein_id', 'protein_id_annotation', 'gene_id'
```

In [7]:

```
print("Is proteodata: ", is_proteodata(adata))
```

```
Is proteodata:  (True, 'peptide')
```

In [8]:

```
adata.obs.head(n=10)
```

Out[8]:

|  | sample\_id | tissue | mouse\_id |
| --- | --- | --- | --- |
| 101\_Brain | 101\_Brain | Brain | 101 |
| 45\_Brain | 45\_Brain | Brain | 45 |
| 66\_Brain | 66\_Brain | Brain | 66 |
| 68\_Brain | 68\_Brain | Brain | 68 |
| 73\_Brain | 73\_Brain | Brain | 73 |
| 80\_Brain | 80\_Brain | Brain | 80 |
| BAT\_101 | BAT\_101 | BAT | 101 |
| BAT\_45 | BAT\_45 | BAT | 45 |
| BAT\_66 | BAT\_66 | BAT | 66 |
| BAT\_68 | BAT\_68 | BAT | 68 |

In [9]:

```
adata.var.head(n=10)
```

Out[9]:

|  | peptide\_id | protein\_id | protein\_id\_annotation | gene\_id |
| --- | --- | --- | --- | --- |
| AAAAAAAAAAAAAAAGAAGK | AAAAAAAAAAAAAAAGAAGK | P55012 | P55012 | Slc12a2 |
| AAAAADLANR | AAAAADLANR | Q9JHS4 | Q9JHS4 | Clpx |
| AAAADGEPLHNEEER | AAAADGEPLHNEEER | Q80WW9 | Q80WW9 | Ddrgk1 |
| AAAAEGARPLER | AAAAEGARPLER | Q80UM7 | Q80UM7 | Mogs |
| AAAAKEEAPK | AAAAKEEAPK | Q91XV3 | Q91XV3 | Basp1 |
| AAAANLC(UniMod:4)PGDVILAIDGFGTESMTHADAQDR | AAAANLC(UniMod:4)PGDVILAIDGFGTESMTHADAQDR | O70209 | O70209 | Pdlim3 |
| AAAAYALGR | AAAAYALGR | Q6ZQ73 | Q6ZQ73 | Cand2 |
| AAADLM(UniMod:35)AYC(UniMod:4)EAHAKEDPLLTPVPASENPFR | AAADLM(UniMod:35)AYC(UniMod:4)EAHAKEDPLLTPVPAS... | P63213 | P63213 | Gng2 |
| AAADLMAYC(UniMod:4)EAHAK | AAADLMAYC(UniMod:4)EAHAK | P63213 | P63213 | Gng2 |
| AAADLMAYC(UniMod:4)EAHAKEDPLLTPVPASENPFR | AAADLMAYC(UniMod:4)EAHAKEDPLLTPVPASENPFR | P63213 | P63213 | Gng2 |

You can define consistent color schemes and category orders for the different tissues and BXD mice used for all figures created by ProteoPy.

In [10]:

```
# Tissues
n_tissues = adata.obs["tissue"].nunique()
cmap = mpl.colormaps["Set2"]
adata.uns["colors_tissue"] = cmap(range(n_tissues)).tolist()

adata.uns["order_tissue"] = ["Brain", "BAT", "Heart", "Liver", "Quad"]

# BXD mice
n_mice = adata.obs["mouse_id"].nunique()
cmap = mpl.colormaps["tab10"]
adata.uns["colors_mouse_id"] = cmap(range(n_mice)).tolist()

adata.uns["order_mouse_id"] = sorted(adata.obs["mouse_id"].unique().tolist())
```

Looking at the sample distribution shows successful data import in agreement with the study design. The dataset includes 5 tissues derived from 8 BXD mice:

In [11]:

```
pr.pl.n_samples_per_category(
    adata,
    category_key="tissue",
    order=adata.uns["order_tissue"],
    color_scheme=adata.uns["colors_tissue"],
)
```

In [12]:

```
pr.pl.n_samples_per_category(
    adata,
    category_key="mouse_id",
    order=adata.uns["order_mouse_id"],
    color_scheme=adata.uns["colors_mouse_id"],
)
```

#### Quality control and preprocessing¶

Similar to the original analysis, we remove peptides from the iRT protein, which was used for calibration, and A2ASS6, which has extraordinarily many
peptides and distorts the analysis.

In [13]:

```
irt_mask = (adata.var["protein_id"] == "iRT_protein")
adata = adata[:, ~irt_mask]
```

In [14]:

```
A2ASS6_mask = (adata.var["protein_id"] == "A2ASS6")
print(f"N peptides for protein A2ASS6: {A2ASS6_mask.sum()}")
```

```
N peptides for protein A2ASS6: 919
```

In [15]:

```
adata = adata[:, ~A2ASS6_mask]
```

Next, peptides that differ only by technical modifications or derived from
missed cleavages are summarized by summing up their intensities.

In [16]:

```
pr.pp.summarize_modifications(adata, method="sum", verbose=True)
pr.pp.summarize_overlapping_peptides(adata)
```

```
Stripping modifications: 31760 peptides -> 29862 unique stripped sequences (method='sum').
```

After these preprocessing steps, the dataset includes approximately 25,000
peptides and 3,800 proteins per sample.

In [17]:

```
pr.pl.n_peptides_per_sample(
    adata,
    zero_to_na=True,
    order_by="tissue",
    order=adata.uns["order_tissue"],
    color_scheme=adata.uns["colors_tissue"],
    print_stats=True,
)
```

```
Global:
 mean_count  std_count  median_count  min_count  max_count  mean_pct  std_pct  median_pct  min_pct  max_pct
    25184.2      199.1       25258.5      24623      25418      98.8      0.8        99.1     96.6     99.7

Per tissue:
tissue  mean_count  std_count  median_count  min_count  max_count  mean_pct  std_pct  median_pct  min_pct  max_pct
   BAT     25181.2      191.5       25077.5      24997      25418      98.8      0.8        98.4     98.0     99.7
 Brain     25269.4        7.2       25268.5      25261      25280      99.1      0.0        99.1     99.1     99.1
 Heart     24860.5      110.9       24858.0      24623      24996      97.5      0.4        97.5     96.6     98.0
 Liver     25369.6       11.8       25372.0      25352      25386      99.5      0.0        99.5     99.4     99.6
  Quad     25240.0       27.7       25240.0      25211      25289      99.0      0.1        99.0     98.9     99.2
```

Out[17]:

```
<Axes: ylabel='#'>
```

In [18]:

```
pr.pl.n_proteins_per_sample(
    adata,
    zero_to_na=True,
    order_by="tissue",
    order=adata.uns["order_tissue"],
    color_scheme=adata.uns["colors_tissue"],
    print_stats=True,
)
```

```
Global:
 mean_count  std_count  median_count  min_count  max_count  mean_pct  std_pct  median_pct  min_pct  max_pct
     3808.8        5.8        3811.5       3790       3815      99.8      0.2        99.9     99.3     99.9

Per tissue:
tissue  mean_count  std_count  median_count  min_count  max_count  mean_pct  std_pct  median_pct  min_pct  max_pct
   BAT      3807.1        5.3        3805.0       3801       3814      99.7      0.1        99.7     99.6     99.9
 Brain      3812.1        0.6        3812.0       3811       3813      99.9      0.0        99.9     99.8     99.9
 Heart      3800.1        4.6        3801.0       3790       3806      99.6      0.1        99.6     99.3     99.7
 Liver      3814.0        0.5        3814.0       3813       3815      99.9      0.0        99.9     99.9     99.9
  Quad      3810.8        1.5        3811.0       3808       3813      99.8      0.0        99.8     99.8     99.9
```

Out[18]:

```
<Axes: ylabel='#'>
```

Since proteoform group inference depends on high peptide coverage, we also inspect the distribution of peptides per protein:

In [19]:

```
pr.pl.n_peptides_per_protein(adata, print_stats=True)
```

```
    mean  median  mode  variance  min  max
6.680115     4.0     1 70.811955    1  118
```

Out[19]:

```
<Axes: xlabel='Number of peptide_id per protein_id', ylabel='# protein_id'>
```

Note that no log transformation, data normalization or missing value
imputation is performed here. This is because of the overall very high data
consistency across the dataset and because COPF is based on covariation
rather than direct intensity differences across conditions. This is also in
agreement with the analysis by Bludau et al. (2021). However, other datasets
might require additional preprocessing steps.

#### Exploratory analysis of peptide-level data¶

Before proteoform inference, ProteoPy can be used to perform classical exploratory data analysis steps, for example to investigate the primary sources of variation in the dataset.

##### Sample correlation matrix¶

High intra-tissue correlation and lower inter-tissue correlation show that peptide intensities are strongly driven by the tissue of origin and not by the mouse strain.

In [20]:

```
pr.pl.sample_correlation_matrix(
    adata,
    method="pearson",
    margin_color="tissue",
    color_scheme=adata.uns["colors_tissue"],
)
```

##### Dimensionality reduction¶

The AnnData-based proteodata object created and used by ProteoPy is directly compatible with other AnnData based python packages such as scanpy (`import scanpy as sc`), which can for example be used for dimensionality reduction and clustering.

Here, the PCA and UMAP analyses further confirm that tissue origin is the dominant source of variation in the dataset.

###### PCA analysis¶

In [21]:

```
sc.tl.pca(adata)

with rc_context({"figure.figsize": (5, 3)}):
    sc.pl.pca_variance_ratio(adata, n_pcs=50, log=True)
```

In [22]:

```
sc.pl.pca(
    adata,
    color=["tissue", "tissue", "tissue"],
    dimensions=[(0, 1), (1, 2), (2, 3)],
    ncols=3,
    size=90,
    palette=adata.uns["colors_tissue"],
)

sc.pl.pca(
    adata,
    color=["mouse_id", "mouse_id", "mouse_id"],
    dimensions=[(0, 1), (1, 2), (2, 3)],
    ncols=3,
    size=90,
    palette=adata.uns["colors_mouse_id"],
)
```

###### UMAP¶

In [23]:

```
sc.pp.neighbors(adata, n_neighbors=4)
sc.tl.umap(adata)

sc.pl.umap(
    adata,
    color=["tissue"],
    size=100,
    palette=adata.uns["colors_tissue"],
)

sc.pl.umap(
    adata,
    color=["mouse_id"],
    size=100,
    palette=adata.uns["colors_mouse_id"],
)
```

```
/home/ifichtner/miniforge3/envs/proteopy-0-1-1/lib/python3.10/site-packages/sklearn/manifold/_spectral_embedding.py:328: UserWarning: Graph is not fully connected, spectral embedding may not work as expected.
  warnings.warn(
```

#### Proteoform inference¶

ProteoPy includes the core functionality of the COPF workflow: (1) compute pairwise peptide correlations between all peptides of a protein, (2) perform hierarchical clustering on the correlation distances, (3) cut each dendrogram into two clusters representing proteoform groups, and (4) score the between- vs. within-cluster correlation difference. Proteins require at least 4 peptides for meaningful clustering.

In [24]:

```
# Require ≥4 peptides per protein for COPF analysis
pr.pp.filter_proteins_by_peptide_count(adata, min_count=4)
pr.pp.remove_zero_variance_vars(adata)
```

```
Removed 1770 proteins and 2984 peptides.
```

In [25]:

```
# COPF pipeline: correlate, cluster, score
pr.tl.pairwise_peptide_correlations(adata)
pr.tl.peptide_dendograms_by_correlation(adata, method="agglomerative-hierarchical-clustering")
pr.tl.peptide_clusters_from_dendograms(adata, n_clusters=2, min_peptides_per_cluster=2)
pr.tl.proteoform_scores(adata, min_score=0.1, min_pval_adj=0.1)
```

In [26]:

```
# Remove COPF outlier peptides (cluster_id == 1000000)
copf_outliers = (adata.var["cluster_id"] == 1000000)
adata = adata[:, ~copf_outliers]
```

The pseudo-volcano plot shows proteoform scores vs. adjusted p-values (related to Figure 6B in Bludau et al. (2021)). The two highlighted proteins — Ldb3 (Q9JKS4) and Sorbs2 (Q3UTJ2) — are examined in detail below.

In [27]:

```
bludauI_proteoforms = ["Q9JKS4", "Q3UTJ2"]
pr.pl.proteoform_scores(
    adata,
    adj=True,
    pval_threshold=0.1,
    score_threshold=0.1,
    highlight_prots=bludauI_proteoforms,
)
```

Out[27]:

```
<Axes: xlabel='Proteoform Score', ylabel='adj. p-value'>
```

In [28]:

```
pf_df = pr.get.proteoforms_df(adata, only_proteins=True)
n_proteoforms = pf_df[pf_df["is_proteoform"] == 1].shape[0]
print(f"{n_proteoforms} significant proteoforms inferred.")
```

```
65 significant proteoforms inferred.
```

#### Proteoform exploration¶

ProteoPy also recovered the two exemplary proteins with proteoform groups highlighted in Bludau et al (2021): LIM domain-binding protein 3 (Ldb3, Q9JKS4) and Sorbin and SH3 domain-containing protein 2 (Sorbs2, Q3UTJ2).

##### LIM domain-binding protein 3 — Ldb3 (Q9JKS4)¶

Ldb3 is a muscle-specific protein. COPF assigns its peptides to two tissue-specific proteoform groups, consistent with known alternative splice variants (reproduces Figure 7A).

In [29]:

```
pr.pl.proteoform_intensities(
    adata,
    protein_ids="Q9JKS4",
    order_by="tissue",
    order=adata.uns["order_tissue"],
    xlab_rotation=45,
)
```

In [30]:

```
pr.get.proteoforms_df(adata, proteins="Q9JKS4")
```

Out[30]:

|  | protein\_id | peptide\_id | cluster\_id | proteoform\_score | proteoform\_score\_pval | proteoform\_score\_pval\_adj | is\_proteoform |
| --- | --- | --- | --- | --- | --- | --- | --- |
| 0 | Q9JKS4 | QYNNPIGLYSAETLR | 0.0 | 0.529813 | 0.002771 | 0.062994 | 1.0 |
| 1 | Q9JKS4 | ASSEGAQGSVSPK | 0.0 | 0.529813 | 0.002771 | 0.062994 | 1.0 |
| 2 | Q9JKS4 | VLPGPSQPR | 0.0 | 0.529813 | 0.002771 | 0.062994 | 1.0 |
| 3 | Q9JKS4 | EMAQMYQMSLR | 0.0 | 0.529813 | 0.002771 | 0.062994 | 1.0 |
| 4 | Q9JKS4 | ILAQMTGTEYMQDPDEEALRR | 1.0 | 0.529813 | 0.002771 | 0.062994 | 1.0 |
| 5 | Q9JKS4 | IMGEVMHALR | 1.0 | 0.529813 | 0.002771 | 0.062994 | 1.0 |
| 6 | Q9JKS4 | QTWHTTCFVCAACK | 1.0 | 0.529813 | 0.002771 | 0.062994 | 1.0 |
| 7 | Q9JKS4 | SKRPIPISTTAPPIQSPLPVIPHQK | 1.0 | 0.529813 | 0.002771 | 0.062994 | 1.0 |
| 8 | Q9JKS4 | SASYNLSLTLQK | 1.0 | 0.529813 | 0.002771 | 0.062994 | 1.0 |
| 9 | Q9JKS4 | SWHPEEFNCAYCK | 1.0 | 0.529813 | 0.002771 | 0.062994 | 1.0 |
| 10 | Q9JKS4 | ASGAGLLGGSLPVKDLAVDSASPVYQAVIK | 1.0 | 0.529813 | 0.002771 | 0.062994 | 1.0 |
| 11 | Q9JKS4 | TPLCGHCNNVIR | 1.0 | 0.529813 | 0.002771 | 0.062994 | 1.0 |
| 12 | Q9JKS4 | TQSKPEDEADEWARR | 1.0 | 0.529813 | 0.002771 | 0.062994 | 1.0 |
| 13 | Q9JKS4 | TSLADVCFVEEQNNVYCER | 1.0 | 0.529813 | 0.002771 | 0.062994 | 1.0 |
| 14 | Q9JKS4 | AAQSQLSQGDLVVAIDGVNTDTMTHLEAQNK | 1.0 | 0.529813 | 0.002771 | 0.062994 | 1.0 |
| 15 | Q9JKS4 | CYEQFFAPICAK | 1.0 | 0.529813 | 0.002771 | 0.062994 | 1.0 |
| 16 | Q9JKS4 | LQGGKDFNMPLTISR | 1.0 | 0.529813 | 0.002771 | 0.062994 | 1.0 |
| 17 | Q9JKS4 | GAPAYNPTGPQVTPLAR | 1.0 | 0.529813 | 0.002771 | 0.062994 | 1.0 |
| 18 | Q9JKS4 | GPFLVAMGR | 1.0 | 0.529813 | 0.002771 | 0.062994 | 1.0 |

##### Ldb3 peptide sequence map (reproduces Figure 7C)¶

Mapping detected peptides onto the canonical protein sequence and annotated UniProt alternative sequences confirms that the two proteoform groups correspond to known splice variants.

In [31]:

```
# Canonical sequence and UniProt alternative sequences for Q9JKS4
# Source: https://www.uniprot.org/uniprotkb/Q9JKS4/entry#sequences
seq = "MSYSVTLTGPGPWGFRLQGGKDFNMPLTISRITPGSKAAQSQLSQGDLVVAIDGVNTDTMTHLEAQNKIKSASYNLSLTLQKSKRPIPISTTAPPIQSPLPVIPHQKDPALDTNGSLATPSPSPEARASPGALEFGDTFSSSFSQTSVCSPLMEASGPVLPLGSPVAKASSEGAQGSVSPKVLPGPSQPRQYNNPIGLYSAETLREMAQMYQMSLRGKASGAGLLGGSLPVKDLAVDSASPVYQAVIKTQSKPEDEADEWARRSSNLQSRSFRILAQMTGTEYMQDPDEEALRRSSTPIEHAPVCTSQATSPLLPASAQSPAAASPIAASPTLATAAATHAAAASAAGPAASPVENPRPQASAYSPAAAASPAPSAHTSYSEGPAAPAPKPRVVTTASIRPSVYQPVPASSYSPSPGANYSPTPYTPSPAPAYTPSPAPTYTPSPAPTYSPSPAPAYTPSPAPNYTPTPSAAYSGGPSESASRPPWVTDDSFSQKFAPGKSTTTVSKQTLPRGAPAYNPTGPQVTPLARGTFQRAERFPASSRTPLCGHCNNVIRGPFLVAMGRSWHPEEFNCAYCKTSLADVCFVEEQNNVYCERCYEQFFAPICAKCNTKIMGEVMHALRQTWHTTCFVCAACKKPFGNSLFHMEDGEPYCEKDYINLFSTKCHGCDFPVEAGDKFIEALGHTWHDTCFICAVCHVNLEGQPFYSKKDKPLCKKHAHAINV"
alt_seqs = {
    "Q9JKS4-2_0": {"seq_coord": (107, 227), "group": "alt_2"},
    "Q9JKS4-3_0": {"seq_coord": (295, 357), "group": "alt_3"},
    "Q9JKS4-4_0": {"seq_coord": (107, 227), "group": "alt_4"},
    "Q9JKS4-4_1": {"seq_coord": (295, 357), "group": "alt_4"},
    "Q9JKS4-5_0": {"seq_coord": (295, 327), "group": "alt_5"},
    "Q9JKS4-5_1": {"seq_coord": (328, 723), "group": "alt_5"},
    "Q9JKS4-6_0": {"seq_coord": (107, 227), "group": "alt_6"},
    "Q9JKS4-6_1": {"seq_coord": (295, 327), "group": "alt_6"},
    "Q9JKS4-6_2": {"seq_coord": (328, 723), "group": "alt_6"},
}

pr.pl.peptides_on_prot_sequence(
    adata,
    protein_id="Q9JKS4",
    group_by="proteoform_id",
    ref_sequence=seq,
    add_sequences=alt_seqs,
    title="LIM domain binding protein 3 (Q9JKS4)",
    figsize=(8, 4),
)
```

Out[31]:

```
<Axes: title={'center': 'LIM domain binding protein 3 (Q9JKS4)'}, xlabel='Position'>
```

##### Sorbin and SH3 domain-containing protein 2 — Sorbs2 (Q3UTJ2)¶

Sorbs2 peptides are assigned to two proteoform groups with distinct tissue expression patterns: one group is abundant across brain, heart, and liver, while the other is brain-specific (reproduces Figure 7D).

In [32]:

```
pr.pl.proteoform_intensities(
    adata,
    protein_ids="Q3UTJ2",
    order_by="tissue",
    order=adata.uns["order_tissue"],
    xlab_rotation=45,
)
```

In [33]:

```
pr.get.proteoforms_df(adata, proteins="Q3UTJ2")
```

Out[33]:

|  | protein\_id | peptide\_id | cluster\_id | proteoform\_score | proteoform\_score\_pval | proteoform\_score\_pval\_adj | is\_proteoform |
| --- | --- | --- | --- | --- | --- | --- | --- |
| 0 | Q3UTJ2 | LAFLVSPVPFR | 0.0 | 0.619637 | 0.000343 | 0.01393 | 1.0 |
| 1 | Q3UTJ2 | ASVVEALDSALKDICDQIK | 0.0 | 0.619637 | 0.000343 | 0.01393 | 1.0 |
| 2 | Q3UTJ2 | QGIFPVSYVEVVKR | 1.0 | 0.619637 | 0.000343 | 0.01393 | 1.0 |
| 3 | Q3UTJ2 | APHYPGIGPVDESGIPTAIR | 1.0 | 0.619637 | 0.000343 | 0.01393 | 1.0 |
| 4 | Q3UTJ2 | SFISSSPSSPSR | 1.0 | 0.619637 | 0.000343 | 0.01393 | 1.0 |
| 5 | Q3UTJ2 | SIFEYEPGK | 1.0 | 0.619637 | 0.000343 | 0.01393 | 1.0 |
| 6 | Q3UTJ2 | AQPARPPPPVQPGEIGEAIAK | 1.0 | 0.619637 | 0.000343 | 0.01393 | 1.0 |
| 7 | Q3UTJ2 | SYSSTLTDLGR | 1.0 | 0.619637 | 0.000343 | 0.01393 | 1.0 |
| 8 | Q3UTJ2 | VGIFPISYVEK | 1.0 | 0.619637 | 0.000343 | 0.01393 | 1.0 |
| 9 | Q3UTJ2 | ADLPGSSSTFTK | 1.0 | 0.619637 | 0.000343 | 0.01393 | 1.0 |
| 10 | Q3UTJ2 | GLGDQSSSR | 1.0 | 0.619637 | 0.000343 | 0.01393 | 1.0 |

##### Sorbs2 peptide sequence map (reproduces Figure 7F)¶

The brain-specific proteoform group maps to a region covered by alternative sequences 3, 4 and 5 in UniProt (10 in Bludau et. al. Figure 7F), a brain-specific splice variant.

In [34]:

```
# Canonical sequence and UniProt alternative sequences for Q3UTJ2
# Source: https://www.uniprot.org/uniprotkb/Q3UTJ2/entry#sequences
seq = "MNTDSGGCARKRAAMSVTLTSVKRVQSSPNLLAAGRESQSPDSAWRSYNDRNPETLNGDATYSSLAAKGFRSVRPNLQDKRSPTQSQITINGNSGGAVSPVSYYQRPFSPSAYSLPASLNSSIIMQHGRSLDSAETYSQHAQSLDGTMGSSIPLYRSSEEEKRVTVIKAPHYPGIGPVDESGIPTAIRTTVDRPKDWYKTMFKQIHMVHKPGLYNSPYSAQSHPAAKTQTYRPLSKSHSDNGTDAFKEVPSPVPPPHVPPRPRDQSSTLKHDWDPPDRKVDTRKFRSEPRSIFEYEPGKSSILQHERPVSIYQSSIDRSLERPSSSASMAGDFRKRRKSEPAVGPLRGLGDQSSSRTSPGRADLPGSSSTFTKSFISSSPSSPSRAQGGDDSKMCPPLCSYSGLNGTPSGELECCNAYRQHLDVPGDSQRAITFKNGWQMARQNAEIWSSTEETVSPKIKSRSCDDLLNDDCDSFPDPKTKSESMGSLLCEEDSKESCPMTWASPYIQEVCGNSRSRLKHRSAHNAPGFLKMYKKMHRINRKDLMNSEVICSVKSRILQYEKEQQHRGLLHGWSQSSTEEVPRDVVPTRISEFEKLIQKSKSMPNLGDEMLSPITLEPPQNGLCPKRRFSIESLLEEETQVRHPSQGQRSCKSNTLVPIHIEVTSDEQPRTHMEFSDSDQDGVVSDHSDYVHVEGSSFCSESDFDHFSFTSSESFYGSSHHHHHHHHHHRHLISSCKGRCPASYTRFTTMLKHERAKHENMDRPRRQEMDPGLSKLAFLVSPVPFRRKKILTPQKQTEKAKCKASVVEALDSALKDICDQIKAEKRRGSLPDNSILHRLISELLPQIPERNSSLHALKRSPMHQPFHPLPPDGASHCPLYQNDCGRMPHSASFPDVDTTSNYHAQDYGSALSLQDHESPRSYSSTLTDLGRSASRERRGTPEKEKLPAKAVYDFKAQTSKELSFKKGDTVYILRKIDQNWYEGEHHGRVGIFPISYVEKLTPPEKAQPARPPPPVQPGEIGEAIAKYNFNADTNVELSLRKGDRIILLKRVDQNWYEGKIPGTNRQGIFPVSYVEVVKRNAKGAEDYPDPPLPHSYSSDRIYTLSSNKPQRPGFSHENIQGGGEPFQALYNYTPRNEDELELRESDVVDVMEKCDDGWFVGTSRRTKFFGTFPGNYVKRL"
alt_seqs = {
    "Q3UTJ2-2_0": {"seq_coord": (210, 211), "group": "alt_2"},
    "Q3UTJ2-2_1": {"seq_coord": (307, 308), "group": "alt_2"},
    "Q3UTJ2-3_0": {"seq_coord": (3, 34), "group": "alt_3"},
    "Q3UTJ2-3_1": {"seq_coord": (210, 211), "group": "alt_3"},
    "Q3UTJ2-3_2": {"seq_coord": (387, 914), "group": "alt_3"},
    "Q3UTJ2-3_3": {"seq_coord": (1125, 1180), "group": "alt_3"},
    "Q3UTJ2-4_0": {"seq_coord": (3, 34), "group": "alt_4"},
    "Q3UTJ2-4_1": {"seq_coord": (387, 914), "group": "alt_4"},
    "Q3UTJ2-5_0": {"seq_coord": (44, 67), "group": "alt_5"},
    "Q3UTJ2-5_1": {"seq_coord": (210, 211), "group": "alt_5"},
    "Q3UTJ2-5_2": {"seq_coord": (307, 308), "group": "alt_5"},
    "Q3UTJ2-5_3": {"seq_coord": (387, 914), "group": "alt_5"},
    "Q3UTJ2-5_4": {"seq_coord": (1125, 1135), "group": "alt_5"},
    "Q3UTJ2-6_0": {"seq_coord": (44, 67), "group": "alt_6"},
    "Q3UTJ2-6_1": {"seq_coord": (210, 211), "group": "alt_6"},
    "Q3UTJ2-6_2": {"seq_coord": (308, 316), "group": "alt_6"},
    "Q3UTJ2-6_3": {"seq_coord": (316, 1180), "group": "alt_6"},
    "Q3UTJ2-7_0": {"seq_coord": (310, 313), "group": "alt_7"},
    "Q3UTJ2-7_1": {"seq_coord": (313, 1180), "group": "alt_7"},
}

pr.pl.peptides_on_prot_sequence(
    adata,
    protein_id="Q3UTJ2",
    group_by="proteoform_id",
    ref_sequence=seq,
    add_sequences=alt_seqs,
    title="Sorbin and SH3 domain-containing protein 2 (Q3UTJ2)",
    figsize=(8, 4),
)
```

Out[34]:

```
<Axes: title={'center': 'Sorbin and SH3 domain-containing protein 2 (Q3UTJ2)'}, xlabel='Position'>
```

#### Proteoform quantification and statistical analysis¶

To test if the inferred proteoform groups are indeed tissue-specific, a one-way ANOVA analysis can be performed directly using ProteoPy functions. In strong agreement with Bludau et al. (2021), our analysis revealed that 59 out of 65 proteoform-containing proteins (90.8%) are expressed in a tissue-specific manner.

In [35]:

```
# Aggregate peptide intensities to proteoform level
adata_pfs = adata.copy()
pr.pp.quantify_proteoforms(adata_pfs, group_by="proteoform_id")
```

In [36]:

```
# Retain only significant proteoforms (score >= 0.1, adj. p-value <= 0.1)
pf_mask = (
    (adata_pfs.var["proteoform_score"].astype(float) >= 0.1)
    & (adata_pfs.var["proteoform_score_pval_adj"].astype(float) <= 0.1)
)
adata_pfs = adata_pfs[:, pf_mask].copy()
```

In [37]:

```
# Log2 transform for statistical testing
adata_pfs.layers["raw"] = adata_pfs.X
adata_pfs.X[adata_pfs.X == 0] = np.nan
adata_pfs.X = np.log2(adata_pfs.X)
adata_pfs.X[np.isnan(adata_pfs.X)] = 0
```

In [38]:

```
# One-way ANOVA across tissues
pr.tl.differential_abundance(
    adata_pfs,
    method="anova_oneway",
    multitest_correction="bonferroni",
    group_by="tissue",
    alpha=0.01,
    space="log",
)
```

```
Saved test results in .varm['anova_oneway;tissue;all']
```

In [39]:

```
anova_results = pr.get.differential_abundance_df(adata_pfs, keys="anova_oneway;tissue;all")
anova_results.rename(columns={"var_id": "proteoform_id"}, inplace=True)
anova_results
```

Out[39]:

|  | proteoform\_id | test\_type | group\_by | design | fstat | pval | mean\_Brain | mean\_BAT | mean\_Heart | mean\_Liver | mean\_Quad | pval\_adj | is\_diff\_abundant |
| --- | --- | --- | --- | --- | --- | --- | --- | --- | --- | --- | --- | --- | --- |
| 0 | O08601\_0 | anova\_oneway | tissue | all | 29.998692 | 7.135979e-11 | 15.447875 | 16.438822 | 13.642355 | 15.286296 | 13.226680 | 9.276772e-09 | True |
| 1 | O08601\_1 | anova\_oneway | tissue | all | 309.316738 | 8.846917e-27 | 17.173194 | 16.274997 | 16.297549 | 19.401295 | 16.539176 | 1.150099e-24 | True |
| 2 | O54724\_0 | anova\_oneway | tissue | all | 12.603239 | 1.878084e-06 | 15.879478 | 13.639894 | 8.489357 | 10.545068 | 11.828292 | 2.441509e-04 | True |
| 3 | O54724\_1 | anova\_oneway | tissue | all | 184.250825 | 5.444913e-23 | 16.359031 | 19.356209 | 18.207868 | 15.845380 | 17.226098 | 7.078387e-21 | True |
| 4 | O54749\_0 | anova\_oneway | tissue | all | 5.288221 | 1.938883e-03 | 13.162980 | 12.879554 | 14.428646 | 13.568651 | 13.071253 | 2.520548e-01 | False |
| ... | ... | ... | ... | ... | ... | ... | ... | ... | ... | ... | ... | ... | ... |
| 125 | Q9JKS4\_1 | anova\_oneway | tissue | all | 66.936903 | 6.594055e-16 | 17.255804 | 17.226273 | 19.196674 | 17.040296 | 19.944539 | 8.572271e-14 | True |
| 126 | Q9QYR6\_0 | anova\_oneway | tissue | all | 14.856602 | 3.441306e-07 | 15.506743 | 12.560879 | 13.676294 | 13.230452 | 13.061866 | 4.473698e-05 | True |
| 127 | Q9QYR6\_1 | anova\_oneway | tissue | all | 646.764070 | 2.861598e-32 | 20.504493 | 16.481401 | 16.861090 | 17.344528 | 16.848407 | 3.720078e-30 | True |
| 128 | Q9Z204\_0 | anova\_oneway | tissue | all | 75.275766 | 1.071660e-16 | 16.117296 | 12.403909 | 12.737527 | 13.374945 | 12.716591 | 1.393159e-14 | True |
| 129 | Q9Z204\_1 | anova\_oneway | tissue | all | 152.715690 | 1.224770e-21 | 15.790335 | 14.611674 | 13.502690 | 16.977432 | 13.218752 | 1.592201e-19 | True |

130 rows × 13 columns

In [40]:

```
# Count proteins with all proteoforms being significantly tissue-specific
protein_id_map = adata_pfs.var[["protein_id", "protein_id_old"]].set_index("protein_id")["protein_id_old"]
anova_results["protein_id"] = anova_results["proteoform_id"].map(protein_id_map)
n_tissue_specific_pfs = anova_results.groupby("protein_id")["is_diff_abundant"].all().sum()

print(f"{n_tissue_specific_pfs} tissue-specific proteoform groups found via ANOVA.")
```

```
59 tissue-specific proteoform groups found via ANOVA.
```

#### Summary¶

This notebook reproduced the core proteoform inference workflow from Bludau et al. (2021) using ProteoPy. Key results:

- **Proteoform detection**: COPF identified 65 proteins with proteoform groups, comparable to the 63 reported in the original study (Figure 6B).
- **Biological verification**: The tissue-specific proteoform assignments for Ldb3 (Q9JKS4) and Sorbs2 (Q3UTJ2) match the published findings (Figures 7A, 7C, 7D, 7F), with peptide-to-sequence mappings consistent with known alternative splice variants.
- **Tissue specificity**: ANOVA analysis recovered 59 tissue-specific proteoform groups (90.8% of the inferred proteoform groups) — similar to the 56 tissue-specific proteoform groups (88.9% of the inferred proteoform groups) reported by Bludau et al.

Minor discrepancies between reported numbers are expected due to implementation details in preprocessing (e.g., peptide modification summarization, overlapping peptide handling) and statistical testing.

In [41]:

```
!pip freeze
```

```
adjustText==1.3.0
anndata==0.11.4
anyio==4.12.1
argon2-cffi==25.1.0
argon2-cffi-bindings==25.1.0
array-api-compat==1.14.0
arrow==1.4.0
asttokens==3.0.1
async-lru==2.3.0
attrs==26.1.0
babel==2.18.0
beautifulsoup4==4.14.3
biopython==1.86
bleach==6.3.0
certifi==2026.2.25
cffi==2.0.0
charset-normalizer==3.4.6
comm==0.2.3
contourpy==1.3.2
cycler==0.12.1
debugpy==1.8.20
decorator==5.2.1
defusedxml==0.7.1
et_xmlfile==2.0.0
exceptiongroup==1.3.1
executing==2.2.1
fastjsonschema==2.21.2
fonttools==4.62.1
fqdn==1.5.1
h11==0.16.0
h5py==3.16.0
httpcore==1.0.9
httpx==0.28.1
idna==3.11
igraph==1.0.0
ipykernel==7.2.0
ipython==8.38.0
isoduration==20.11.0
jedi==0.19.2
Jinja2==3.1.6
joblib==1.5.3
json5==0.13.0
jsonpointer==3.1.1
jsonschema==4.26.0
jsonschema-specifications==2025.9.1
jupyter-events==0.12.0
jupyter-lsp==2.3.0
jupyter_client==8.8.0
jupyter_core==5.9.1
jupyter_server==2.17.0
jupyter_server_terminals==0.5.4
jupyterlab==4.5.6
jupyterlab_pygments==0.3.0
jupyterlab_server==2.28.0
kiwisolver==1.5.0
lark==1.3.1
legacy-api-wrap==1.5
llvmlite==0.46.0
MarkupSafe==3.0.3
matplotlib==3.10.8
matplotlib-inline==0.2.1
mistune==3.2.0
natsort==8.4.0
nbclient==0.10.4
nbconvert==7.17.0
nbformat==5.10.4
nest-asyncio==1.6.0
networkx==3.4.2
notebook_shim==0.2.4
numba==0.64.0
numpy==2.2.6
openpyxl==3.1.5
overrides==7.7.0
packaging @ file:///home/conda/feedstock_root/build_artifacts/bld/rattler-build_packaging_1769093650/work
pandas==2.3.3
pandocfilters==1.5.1
parso==0.8.6
patsy==1.0.2
pexpect==4.9.0
pillow==12.1.1
platformdirs==4.9.4
pooch==1.9.0
prometheus_client==0.24.1
prompt_toolkit==3.0.52
proteopy==0.1.1
psutil==7.2.2
ptyprocess==0.7.0
pure_eval==0.2.3
pyarrow==23.0.1
pycparser==3.0
Pygments==2.19.2
pynndescent==0.6.0
pyparsing==3.3.2
python-dateutil==2.9.0.post0
python-igraph==1.0.0
python-json-logger==4.0.0
pytz==2026.1.post1
PyYAML==6.0.3
pyzmq==27.1.0
referencing==0.37.0
requests==2.32.5
rfc3339-validator==0.1.4
rfc3986-validator==0.1.1
rfc3987-syntax==1.1.0
rpds-py==0.30.0
scanpy==1.11.5
scikit-learn==1.7.2
scipy==1.15.3
seaborn==0.13.2
Send2Trash==2.1.0
session-info2==0.4
six==1.17.0
soupsieve==2.8.3
stack-data==0.6.3
statsmodels==0.14.6
terminado==0.18.1
texttable==1.7.0
threadpoolctl==3.6.0
tinycss2==1.4.0
tomli==2.4.0
tornado==6.5.5
tqdm==4.67.3
traitlets==5.14.3
typing_extensions==4.15.0
tzdata==2025.3
umap-learn==0.5.11
uri-template==1.3.0
urllib3==2.6.3
wcwidth==0.6.0
webcolors==25.10.0
webencodings==0.5.1
websocket-client==1.9.0
```
